# ORPER: a workflow for constrained SSU rRNA phylogenies

**DOI:** 10.1101/2021.10.04.463113

**Authors:** Luc Cornet, Anne-Catherine Ahn, Annick Wilmotte, Denis Baurain

## Abstract

The continuous increase of sequenced genomes in public repositories makes the choice of interesting bacterial strains for future sequencing projects evermore complicated, as it is difficult to estimate the redundancy between these strains and the already available genomes. Therefore, we developed the Nextflow workflow “ORPER” (containerized in Singularity), which allows determining the phylogenetic position of a collection of organisms in the genomic landscape. ORPER constrains the phylogenetic placement of SSU (16S) rRNA sequences in a multilocus reference tree based on ribosomal protein genes extracted from public genomes. We demonstrate the utility of ORPER on the Cyanobacteria phylum, by placing 152 strains of the BCCM/ULC collection.

## 1. Introduction

Cyanobacteria form a phylum of bacteria having colonized very diversified ecosystems [1]. They are the only bacteria able to perform oxygenic photosynthesis, which appeared at least 2.4 billion years ago [2]. By increasing free atmospheric oxygen, Cyanobacteria had a critical impact on shaping life on Earth [3-4]. Beyond their ecological importance, this phylum also has an evolutionary interest, owing to their key role in the emergence of eukaryotic algae (Archaeplastida) through the primary endosymbiosis having given rise to the plastid [5]. Although the exact mechanisms, which include the generally accepted unicity of the event, remain to be fully understood, it is well known that Cyanobacteria have played a major role in the spread of oxygenic photosynthesis [6]. More recently, the group attracted an additional interest after uncovering, through metagenomic studies, the existence of non-photosynthetic “cyanobacteria”, which were classified under the novel class of Melainabacteria [7].

Due to this importance, published cyanobacterial phylogenies are numerous (see for instance: [8-14]). The number of available genomes has logically followed this interest, rising from a few hundreds in 2013, when Shih et al. [15] improved the coverage of the phylum, to more than 3000 nowadays, according to GenBank statistics. Recent studies have nevertheless demonstrated that cyanobacterial diversity, both for photosynthetic [16] and non-photosynthetic [14] representatives (when considering Melainabacteria as part of Cyanobacteria), are not well covered by the sequencing effort.

The gold standard for the estimation of bacterial diversity remains the SSU (16S) rRNA gene of the small subunit of the ribosomal RNA [17]. This locus is frequently used by scientists and culture collections to evaluate the genomic potential of newly isolated organisms. However, due to the constant and rapid growth of genome repositories, it is difficult for researchers to estimate the redundancy between these public data sources and their own collections of organisms. Here, we release ORPER, standing for “ORganism PlacER”, an automated workflow intended to determine the phylogenetic position of organisms for which only the SSU (16S) rRNA has been determined in the public genomic landscape.

## 2. Methods

### 2.1 Functional overview

The principle of ORPER is to provide an overview of the sequenced coverage (i.e., the diversity of available genomes) of a given taxon and to place SSU (16S) rRNA sequences in this diversity. ORPER first downloads complete genomes of the taxon of interest, then extracts their ribosomal proteins to compute a reference phylogenetic tree, and finally uses this tree to constrain the backbone of a SSU (16S) rRNA phylogeny including the additional strains. The workflow uses two groups: (i) the main group corresponding to the taxonomic group of the SSU (16S) rRNA sequences (the taxon of interest) and (ii) the outgroup to root the phylogenetic tree. The main group will be used to compute a phylogenetic tree to guide the placement of SSU (16S) rRNA sequences; it is therefore named “reference group” in the remaining of this manuscript (**Figure 1**). All steps are embedded in a Nextflow script [18], and a Singularity definition file is provided for containerization [19]. ORPER is available at https://github.com/Lcornet/ORPER.

**Figure 1:**
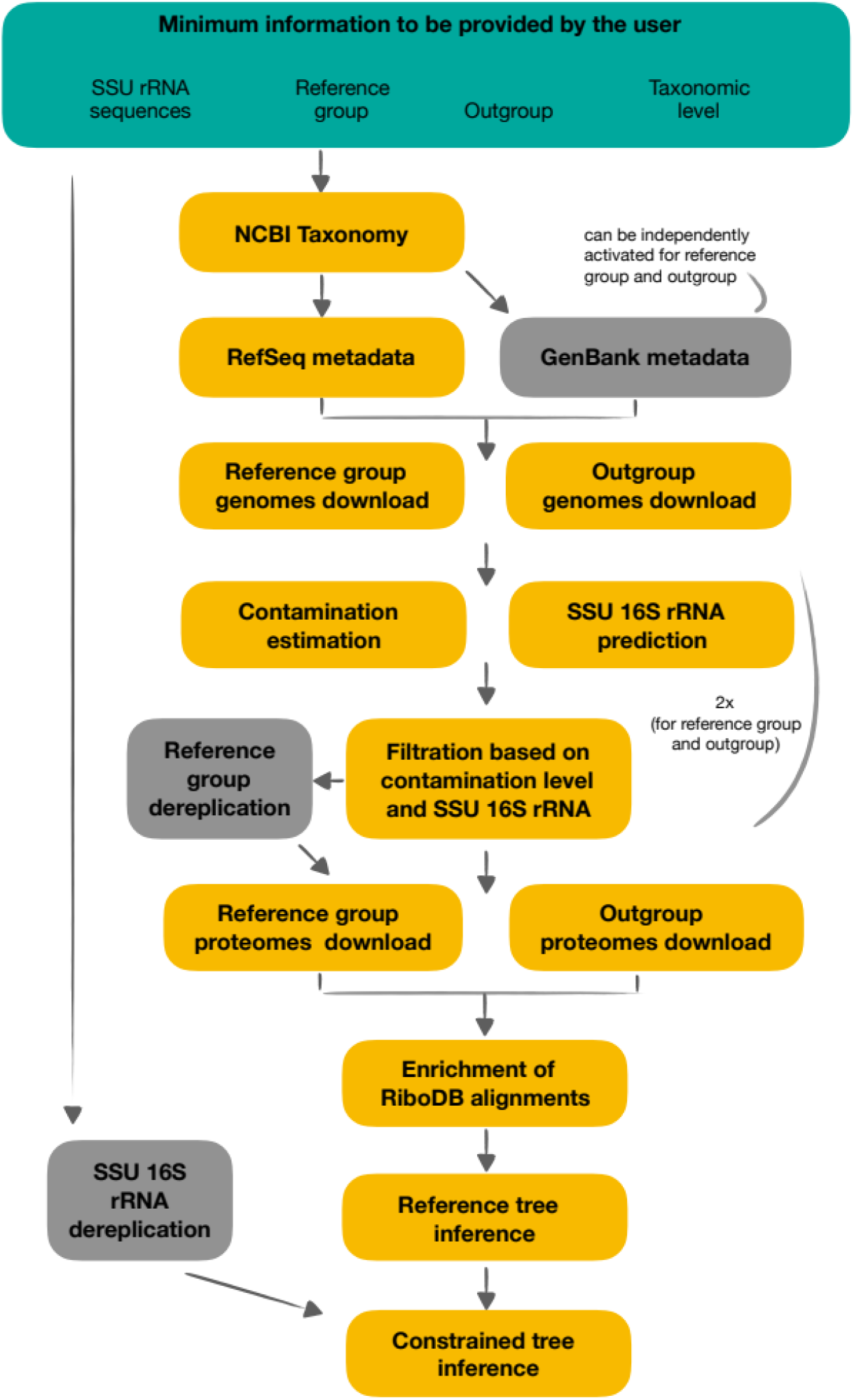
Overview of ORPER workflow. Users should specify at least four pieces of information to run ORPER: (i) their SSU (16S) rRNA sequences, (ii) the taxon of interest, (iii) the outgroup of the phylogeny and (iv) the taxonomic level (Green part). Yellow boxes are mandatory steps of OR-PER whereas grey boxes are optional steps.

### 2.2 Workflow details

#### 2.2.1 Taxonomy and metadata download

ORPER begins by creating a local copy of the National Center for Biotechnology Information (NCBI) Taxonomy [20] with the script *setup-taxdir.pl* v0.211470 from Bio::MUST::Core (D. Baurain; https://metacpan.org/dist/Bio-MUST-Core). Genome accession numbers (i.e., GCF numbers) are fetched from the NCBI Reference Sequence project (RefSeq) [21] and the taxonomy of the corresponding organisms is determined with the script *fetch-tax.pl* v0.211470 (Bio::MUST::Core package). If required, GenBank genomes [22] can be used in the same way for both reference group and outgroup creation, independently. Four taxonomy levels are available in ORPER (phylum, class, order, family) and the user must specify the reference group and the outgroup, separately (**Figure 1**).

#### 2.2.2 Genome filtration and dereplication

*CheckM* v1.1.3 with the “lineage_wf” option, is used to estimate completeness and contamination of the assemblies [23]. Barrnap v0.9, with default options, is used to predict rRNA genes in downloaded genomes (available at https://github.com/tseemann/barrnap). Genomes with a completeness level above 90%, a contamination level below 5%, and at least one predicted SSU (16S) rRNA sequence are retained. A dereplication step of the genomes from the reference group can be optionally carried out using *dRep* [24] and default parameters. *Prodigal*, with default options, is used to obtain conceptual proteomes [25]. All genomes from the reference group that remain after the filtration steps are used whereas only the ten first genomes of the outgroup are used for de novo protein prediction (**Figure 1**).

#### 2.2.3 Reference phylogeny inference

Prokaryotic ribosomal protein alignments from the RiboDB database [26] are downloaded by ORPER once at the first usage. An orthologous enrichment of these alignments with sequences from the remaining proteomes (post-filtration and dereplication) is performed by *Forty-Two v0.210570* [27,28]. These sequences are then aligned using *MUSCLE* v3.8.31 [29] in order to generate new alignment files with only the sequences from the reference group and the outgroup. Conserved sites are selected using *BMGE* v1.12 [30] with moderately severe settings (*entropy cut-off* = 0.5, *gap cut-off* = 0.2). A supermatrix is then generated using *SCaFoS* v1.30k [31] with default settings. Finally, a reference phylogenomic analysis is inferred using *RAxML* v8.2.12 [32] with 100 bootstrap replicates under the PROTGAMMALGF model.

#### 2.2.4 Constrained SSU 16S rRNA phylogeny

The SSU (16S) rRNA sequences provided by the user can be optionally dereplicated using *CD-HIT-EST* v4.8.1 with default parameters [33]. The SSU (16S) rRNA phylogenetic tree is inferred from both the sequences provided by the user and those extracted from the complete genomes using *RAxML* v8.2.12 [32] with 100 bootstrap replicates under the GTRGAMMA model and the phylogenomic tree as a constraint.

### 2.3. Design considerations

ORPER compensates for the lack of phylogenetic resolution of SSU (16S) rRNA gene sequences by using ribosomal protein genes from publicly available genomes to infer a reference multilocus tree, which is then used to constrain the SSU (16S) rRNA phylogeny. It is indeed well known that SSU 16S rRNA suffers, as all single-gene phylogenies, from a lack of phylogenetic resolution [34-38]. Ribosomal protein genes are frequently used to perform phylogenetic placement; for instance, CheckM uses this approach to place genomes before performing the contamination estimation [23].

The NCBI databases, regularly synchronized with the European Nucleotide Archive (ENA) [39], are the most complete public databases. By default, ORPER uses only RefSeq because the latter contains only high-quality genomes [21]. Nevertheless, it might be necessary to use more genomes to estimate the actual sequence coverage of a taxon. It is especially true with metagenomic data that are by design not included in RefSeq [21]. That is why GenBank can be enabled as an option in ORPER. In any case, starting from a NCBI database entails the use of thousands of genomes, which can dramatically increase the computing time. For this reason, we have implemented the optional use of dRep to dereplicate the genomes [24]. This allows the user to decrease the number of genomes while conserving the sequenced diversity [24]. However, this option should be used carefully because the need for dereplication (or not) is dependent on the biological question [40]. For example, the genomic comparison of closely related strains requires using as many genomes as possible to catch individual differences. Finally, genomes in public repositories are not devoid of contamination (i.e., the inclusion of foreign DNA in the genomic data) [41,42]. Therefore, we used CheckM [23], the most commonly used tool for genomic contamination detection, and thresholds from the Genomic Standards Consortium [43] (completeness above 90% and contamination below 5%) to filter our genomes, which is a mandatory step in the workflow.

Nextflow is the latest workflow system. It has been developed to increase reproducibility in science [18]. Nextflow further presents the advantage of exploiting Singularity containers as an operating system [18], which ensures the sustainability of future analyses. Singularity containers [19] correct the security issues of older container systems, thereby offering the possibility to deploy them on HPC systems where security is often an important concern. Owing to these advantages, we chose the combination Nextflow-Singularity for ORPER. Albeit ORPER is a workflow, we designed it as a program with a single command-line interface. The installation of ORPER only requires two shell commands (see https://github.com/Lcornet/ORPER). Moreover, the analysis reported in this study can be replayed with a single command in less than one day using 30 CPU cores (Intel Xeon E5-2640 v4 series) (see https://github.com/Lcornet/ORPER).

## 3. Results and discussion

### 3.1 Case study: BCCM/ULC Cyanobacteria collection

Phylogenomic studies of Cyanobacteria are numerous, notably focussing on the emergence of multicellularity [44-46], the appearance of oxygenic photosynthesis [47-49], or the origin of plastids [9,50-52]. The latter topic is especially controversial [6] with potential origins of plastids either among heterocyst-forming cyanobacteria [2,53] or more early-diverging lineages [9,50-52]. The selection of Cyanobacteria for future sequencing projects thus remains an important issue.

We tested ORPER on this phylum using 152 SSU (16S) rRNA sequences from the BCCM/ULC collection. RefSeq genomes for the “Cyanobacteria” phylum were specified as the reference group, whereas genomes for the “Melainabacteria” phylum available in GenBank were used as the outgroup (**Figure 2**). Dereplication for reference genomes and for SSU (16S) rRNA sequences were both activated. The reference tree inferred by ORPER was based on a supermatrix of 372 organisms x 6246 unambiguously aligned amino-acid positions (7.92% missing character states). The 152 SSU (16S) rRNA input sequences used in this study were dereplicated to 140 sequences at a 95% identity threshold, and were then used to compute the constrained tree (**Table S1**).

**Figure 2:**
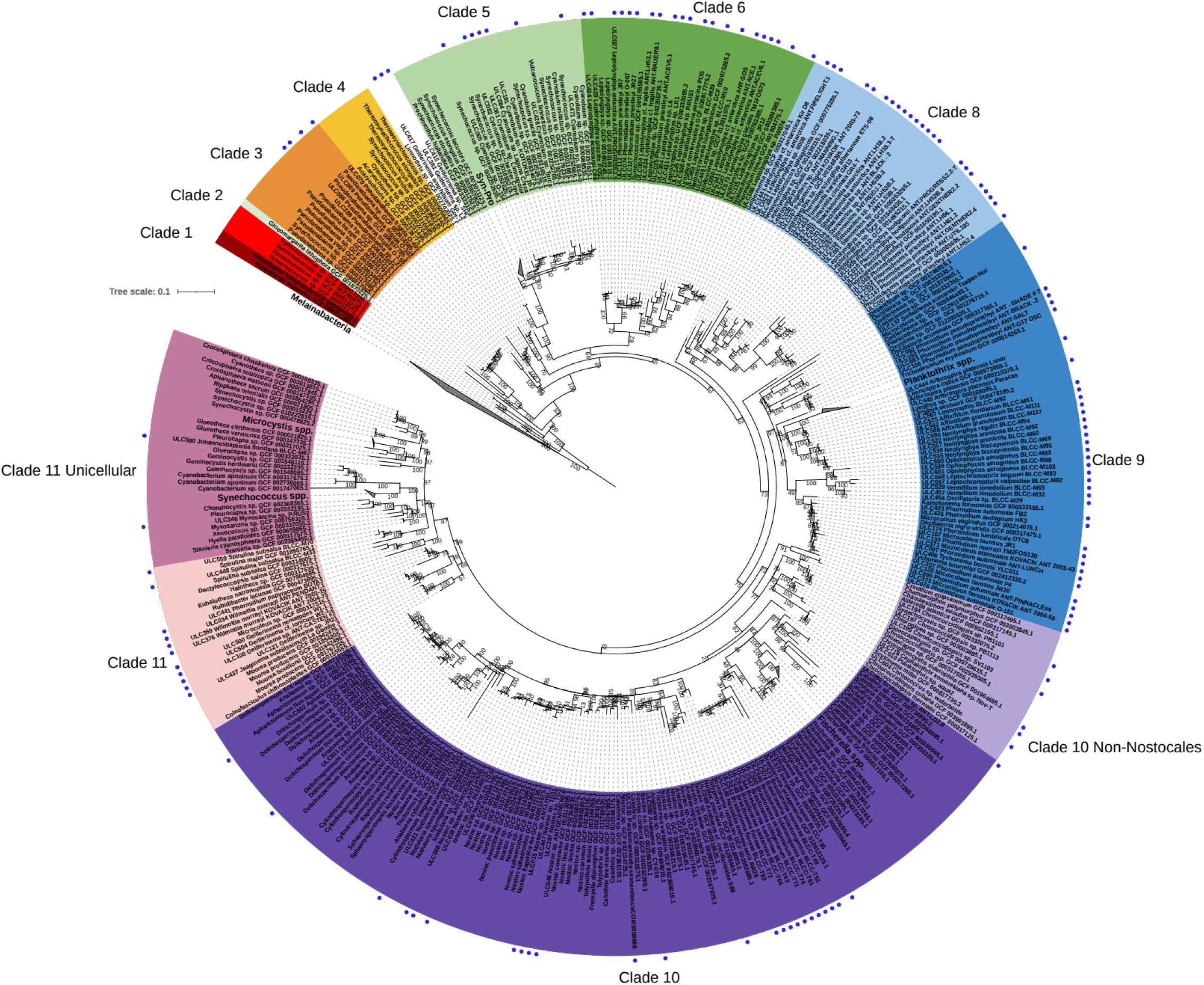
Constrained cyanobacterial phylogenetic tree of the BCCM/ULC collection. The tree is the output of ORPER, a Maximum-likelihood constrained inference computed under the GTRGAMMA model. Clades correspond to the groups defined in Moore et al. (2019) [9]. Clades 10 and 11 have been divided into two sub-clades, adding respectively “Non-Nostocales” and “Unicellular” sub-clades to Moore et al.’s phylogeny. Blue dots indicate ULC/BCCM strains.

We chose to compare the phylogeny inferred by ORPER to the latest multilocus (ribosomal) phylogeny of the cyanobacterial phylum published by Moore et al. (2019), who identified the earliest potential basal position of the plastids [9]. The constrained tree computed by ORPER is comparable to the tree of Moore et al. (2019), with ten out of eleven clades recovered by ORPER (**Figure 2**). The only missing clade, clade 7, was represented by genomes not present in RefSeq [9] or in ULC strains, and is thus logically absent from our phylogeny. The 140 BCCM/ULC strains obtained after dereplication covered the whole diversity of publicly available cyanobacterial genomes. Three BCCM/ULC strains (ULC415, ULC417, ULC381) form a basal clade clustered with *Limnothrix* sp. GCF_002742025.1, which was not present in Moore et al.’s analysis. These three strains would therefore be of high interest for genome sequencing, especially in the context of plastid emergence. Here, we analyzed the cyanobacterial phylum, but ORPER can be used on any bacterial taxon of the NCBI.

## 4. Conclusions

ORPER is a state-of-the-art tool, designed for the phylogenetic placement of SSU (16S) rRNA sequences in phylogenetic tree constrained by a multilocus tree. We demonstrated the utility of ORPER on Cyanobacteria, using sequences from the BCCM/ULC collection, to estimate the phylogenetic position of SSU 16S rRNA sequences among the landscape of sequenced genomes. Its easy-to-use installation process and Singularity containerization makes ORPER a useful tool for culture collections and scientists in their future selection of genomes to sequence.

## Supplementary Materials

The following are available online at www.mdpi.com/xxx/s1, Supplemental file 1: Information on the 152 ULC strains used in this study.

## Author Contributions

LC, DB conceived the study. LC performed all the analyses and drew the figures. AA provided the SSU 16S rRNA gene sequences of the BCCM/ULC strains and their information. LC and DB drafted the manuscript, and AA and AW provided critical reviews of the manuscript. All authors approved the final manuscript.

## Data Availability Statement

ORPER is freely available at https://github.com/Lcornet/ORPER.

## Acknowledgments

The authors thank David Colignon (ULiège) and the CÉCI for their help with computing cluster usage.

