## Supplementary material for "ORPER: a workflow for constrained SSU rRNA phylogenies": Supp file 1

### Supplemental file 1

| Strain accronym | Taxonomic name | Geographical origin | Isolation habitat | Recommended growth temperature (°C) | Recommended growth medium | Reference |
|---|---|---|---|---|---|---|
| ULC001 | *Stenomitos* sp. ANT.L53B.1 | Larsemann Hills, East Antarctica, Prydz Bay, Antarctica | Lake 53, microbial mat | 18 | BG11 | Taton et al., 2011 |
| ULC004 | *Leptolyngbya cf. fragilis* ANT.LH52.1 | Larsemann Hills, East Antarctica, Prydz Bay, Antarctica | Lake 52, microbial mat | 18 | BG11 | Taton et al., 2011 |
| ULC007 | *Phormidesmis priestleyi* ANT.LH52.4 | Larsemann Hills, East Antarctica, Prydz Bay, Antarctica | Lake 52, microbial mat | 18 | BG11 | Taton et al., 2011 |
| ULC008 | *Nostoc* sp. ANT.PROGRESS.2.1 | Larsemann Hills, East Antarctica, Prydz Bay, Antarctica | Lake Progress, microbial mat | 12 | BG110 | |
| ULC009 | *Plectolyngbya hodgsonii* ANT.PROGRESS2.2 T | Larsemann Hills, East Antarctica, Prydz Bay, Antarctica | Lake Progress, microbial mat | 18 | BG11 | Taton et al., 2011 |
| ULC012 | *Plectolyngbya hodgsonii* ANT.GENTNER2.2 | Larsemann Hills, East Antarctica, Prydz Bay, Antarctica | Lake Gentner, microbial mat | 18 | BG11 | Taton et al., 2011 |
| ULC015 | *Nostoc* sp. ANT.LH61.1 | Lake 61, Eastern Antarctica, Larsemann Hills, Antarctica | Lake 61, microbial mat | 12 | BG110 | |
| ULC018 | *Stenomitos* sp. ANT.LH53B.2 | Larsemann Hills, East Antarctica, Prydz Bay, Antarctica | Lake 53, microbial mat | 18 | BG11 | Taton et al., 2011 |
| ULC020 | *Pegethrix* sp. ANT.MANNING.1 | Larsemann Hills, East Antarctica, Prydz Bay, Antarctica | Lake Manning, microbial mat | 18 | BG11 | Taton et al., 2011 |
| ULC021 | *Phormidesmis priestleyi* ANT.GENTNER2.4 | Larsemann Hills, East Antarctica, Prydz Bay, Antarctica | Lake Gentner, microbial mat | 18 | BG11 | Taton et al., 2011 |
| ULC022 | *Phormidesmis priestleyi* ANT.LH61.2 | Larsemann Hills, East Antarctica, Prydz Bay, Antarctica | Lake 61, microbial mat | 18 | BG11 | Taton et al., 2011 |
| ULC023 | *Leptolyngbya cf. antarctica* ANT.FIRELIGHT.1 | Larsemann Hills, East Antarctica, Prydz Bay, Antarctica | Lake Firelight, microbial mat | 12 | BG11 | Taton et al., 2011 |
| ULC024 | *Leptolyngbya* cf. *fragilis* ANT.RAUER8.1 | Rauer Island, East Antarctica, Prydz Bay, Antarctica | Lake Rauer 8, microbial mat | 18 | BG11 | Taton et al., 2011 |
| ULC025 | *Pegethrix* sp. ANT.LH70.1 | Larsemann Hills, East Antarctica, Prydz Bay, Antarctica | Lake 70, microbial mat | 18 | BG11 | Taton et al., 2011 |
| ULC027 | *Leptolyngbya antarctica* ANT.PROGRESS2.5 | Larsemann Hills, East Antarctica, Prydz Bay, Antarctica | Lake Progress, microbial mat | 18 | BG11 | Taton et al., 2011 |

| | | | | | | |
|---|---|---|---|---|---|---|
| ULC029 | *Stenomitos* sp. ANT.LH52B.3 | Larsemann Hills, East Antarctica, Prydz Bay, Antarctica | Lake 52, microbial mat | 12 | BG11 | Taton et al., 2011 |
| ULC030 | *Plectolyngbya hodgsonii* ANT.LH52B.4 | Larsemann Hills, East Antarctica, Prydz Bay, Antarctica | Lake 52, microbial mat | 18 | BG11 | Taton et al., 2011 |
| ULC031 | *Shackletoniella antarctica* ANT.LH18.1(T) | Larsemann Hills, East Antarctica, Prydz Bay, Antarctica | Lake 18, microbial mat | 18 | BG11 | Taton et al., 2011 |
| ULC032 | *Shackletoniella antarctica* ANT.GENTNER2.5 | Larsemann Hills, East Antarctica, Prydz Bay, Antarctica | Lake Gentner, microbial mat | 18 | BG11 | Taton et al., 2011 |
| ULC034 | *Wilmottia murrayi* ANT.PENDANT.1 | Vestfold Hills, East Antarctica, Prydz Bay, Antarctica | Lake Pendant, microbial mat | 18 | BG11 | Taton et al., 2011 |
| ULC035 | *Phormidium priestleyi* ANT.ACEV5.1 | Vestfold Hills, East Antarctica, Prydz Bay, Antarctica | Ace Lake, microbial mat | 12 | BG11 | Taton et al., 2011 |
| ULC036 | *Shackletoniella antarctica* ANT.WATTS.1 | Vestfold Hills, East Antarctica, Prydz Bay, Antarctica | Lake Watts, microbial mat | 18 | BG11 | Taton et al., 2011 |
| ULC037 | *Shackletoniella antarctica* ANT.LH18.2$^{T}$ | Larsemann Hills, East Antarctica, Prydz Bay, Antarctica | Lake 18, microbial mat | 12 | BG11 | Taton et al., 2011; Strunecky et al., 2020 |
| ULC038 | *Nostoc* sp. ANT.LH52B.5 | Larsemann Hills, East Antarctica, Prydz Bay, Antarctica | Lake 52, microbial mat | 18 | BG110 | |
| ULC041 | *Leptolyngbya* cf. *antarctica* ANT.ACE.1 | Vestfold Hills, East Antarctica, Prydz Bay, Antarctica | Ace Lake, microbial mat | 18 | BG11 | Taton et al., 2011 |
| ULC046 | *Nostoc* sp. ANT.GENTNER2.6 | Larsemann Hills, Broknes Peninsula, East Antarctica, Prydz Bay, Antarctica | Lake Gentner 2, microbial mat | 12 | BG110 | |
| ULC047 | *Leptolyngbya* cf. *antarctica* ANT.ACEV6.1 | Vestfold Hills, East Antarctica, Prydz Bay, Antarctica | Ace Lake, microbial mat | 18 | BG11 | Taton et al., 2011 |
| ULC049 | *Phormidesmis priestleyi* ANT.LH66.1 | Larsemann Hills, East Antarctica, Prydz Bay, Antarctica | Lake 66, microbial mat | 18 | BG11 | Taton et al., 2011 |
| ULC057 | *Stenomitos* ANT.REIDJ.1 | Larsemann Hills, East Antarctica, Prydz Bay, Antarctica | Lake Reid, microbial mat | 18 | BG11 | Taton et al., 2011 |
| ULC060 | *Scytonema* sp. ANT.LG2.8 | Larsemann Hills, East Antarctica, Prydz Bay, Antarctica | Lake Gentner, microbial mat | 12 | BG11 | Taton et al., 2011 |
| ULC065 | *Cyanobium* sp. O-154 | Bylot Island, Arctic, Canada | Unknown | 12 | BG11 | |
| ULC066 | *Pseudanabaena frigida* O-155 | Bylot Island, Arctic, Canada | Unknown | 12 | BG11 | |
| ULC069 | *Pseudanabaena frigida* O-302 | Québec, Québec, Sub-arctic, Canada | Lake with clear water | 12 | BG11 | |
| ULC070 | *Pseudanabaena frigida* O-401 | Québec, Québec, Sub-arctic, Canada | Lake with clear water | 12 | BG11 | |
| ULC073 | *Leptolyngbya* | Dufek Massif, | Bottom brine | 12 | BG11 | Fernandez |

| | | | | | | |
|---|---|---|---|---|---|---|
| | *glacialis* TM1FOS73 | Transantarctic Mountains, Antarctica | of Forlidas Pond | | | -Carazo et al., 2011 |
| ULC074 | *Phormidium priestleyi* O-067 | Mc Murdo Ice Shelf, South Victoria Land, Antarctica | Unknown | 12 | BG11 | |
| ULC076 | *Phormidium autumnale* O-151 | Bylot Island, Arctic, Canada | Unknown | 12 | BG11 | |
| ULC080 | *Anabaena* sp. CY-036 | Unknown, New Zealand | Unknown | 12 | BG110 | |
| ULC081 | *Cyanobiu*m sp. Limnopolar | Limnopolar Lake, South Shetland islands, Livingston Island, Antarctica | Unknown | 12 | BG11 | |
| ULC082 | *Cyanobium* sp.Chester Cone | Chester Cone, South Shetland islands, Livingston Island, Antarctica | Unknown | 12 | BG11 | |
| ULC084 | *Cyanobium* sp. Laguna Chica | Laguna Chica, South Shetland islands, Livingston Island, Antarctica | Unknown | 12 | BG11 | |
| ULC088 | *Anabaena* cf. *oscillarioides* S88 | Unknown | Unknown | 12 | BG110 | |
| ULC090 | *Leptolyngbya* cf. *antarctica* ANT-SOS | Mc Murdo Ice Shelf, South Victoria Land, Bratina Island, Antarctica | Son of salt (SOS) pond | 12 | BG11 | Nadeau et al., 2001 |
| ULC092 | *Phormidium pseudopriestleyi* ANT - SHADE # 7 | Mc Murdo Ice Shelf, South Victoria Land, Bratina Island, Antarctica | Shading pond | 12 | BG11 | Nadeau et al., 2001 |
| ULC093 | *Phormidium pseudopriestleyi* ANT-SALT | Mc Murdo Ice Shelf, South Victoria Land, Bratina Island, Antarctica | Salt pond | 12 | BG11 | |
| ULC095 | *Phormidium autumnale* ANT-PINNACLE#4 | Mc Murdo Ice Shelf, South Victoria Land, Bratina Island, Antarctica | Pinnacle pond | 12 | BG11 | |
| ULC097 | *Phormidium autumnale* ANT-LUNCH | Mc Murdo Ice Shelf, South Victoria Land, Bratina island, Antarctica | Pond lunch | 12 | BG11 | Nadeau et al., 2001 |
| ULC100 | *Geitlerinema* sp. ANT-CASTEN #2 | Mc Murdo Ice Shelf, South Victoria Land, Bratina Island, Antarctica | Unknown | 12 | BG11 | |
| ULC102 | *Phormidium pseudopriestleyi* ANT-BRACK -2 | Mc Murdo Ice Shelf, South Victoria Land, Bratina Island, Antarctica | Brack Pond | 12 | BG11 | Nadeau et al., 2001 |
| ULC104 | *Shackletoniella* sp. ANT-BLACK - 2 | Mc Murdo Ice Shelf, South Victoria Land, Bratina Island, Antarctica | Black pond | 12 | BG11 | |
| ULC106 | *Phormidium pseudopriestley*i ANT-G17_OSC | Mc Murdo Ice shelf, South Victoria Land, Bratina Island, Antarctica | Saline pond | 12 | BG11 | |
| ULC107 | *Microcoleus favosus* JR1 | Komarek's seepages, Antarctic Peninsula, James Ross Island, Antarctica | Brown mats in spring area, Slope Creek | 12 | BG11 | Strunecký et al. 2013 |
| ULC110 | *Microcoleus* sp. JR5 | Komarek's seepages, Antarctic Peninsula, | Black biofilm on rocks | 18 | BG11 | Strunecký et al. 2013 |

| | | | | | | |
|---|---|---|---|---|---|---|
| | | James Ross Island, Antarctica | | | | |
| ULC112 | *Phormidium priestleyi* JR7 | Green Lake, Antarctic Peninsula, James Ross Island, Antarctica | Periphyton in littoral of Green Lake, dark-green | 18 | BG11 | |
| ULC115 | *Leptolyngbya* sp. JR12 | Unknown, Antarctic Peninsula, James Ross Island, Antarctica | Unknown | 18 | BG11 | |
| ULC120 | *Microcoleus favosus* JR20 | Antarctic Peninsula, James Ross Island, Antarctica | Rocks wetted by seep | 18 | BG11 | Strunecký et al. 2013 |
| ULC121 | *Geitlerinema* sp. JR21 | Antarctic Peninsula, James Ross Island, Antarctica | Unknown | 12 | BG11 | |
| ULC127 | *Phormidium priestleyi* JR27 | Antarctic Peninsula, James Ross Island, Antarctica | Unknown | 12 | BG11 | |
| ULC130 | *Phormidium murrayi* TM2FOS130 | Forlidas pond, Transantarctic Mountains, Dufek massif, Antarctica | Microbial mat of the littoral zone | 12 | BG11 | Vintila et al., 2011 |
| ULC137 | *Sodalinema komarekii* Tsagan-Nur | Tsagan-Nur lake, Siberia, Russian Federation | Biofilm on the littoral of the lake | 20-25 | Zarrouk+ | Cellamare et al. 2018 |
| ULC138 | *Sodalinema komarekii* Borzinskoye | Siberia, Russian Federation | Lake Borzinskoye | 20-25 | Zarrouk+ | Cellamare et al. 2018 |
| ULC139 | *Sodalinema komarekii* Khil 10-07 | Transbaikalian area, Buryat Republic, Siberia, Russian Federation | Lake Khilganta | 18 | Zarrouk | Cellamare et al. 2018 |
| ULC144 | *Leptolyngbya* cf. *antarctica* Kir D8 | Southern pa20-25 of Buryat Republic, Siberia, Buryat Republic, Russian Federation | Lake Kiranskoye | 20-25 | Zarrouk | |
| ULC146 | *Nostoc* sp. ANT.UTS. 183 | Utsteinen ridge (parcel 21), Dronning Maud land, Sør Rondane Mountains, Antarctica | Black microbial mats and gravel | 18 | BG110 | Fernández -Carazo et al. 2012 |
| ULC147 | *Phormidesmis priestleyi* ANT.UTS.195 | Utsteinen Nunatak, Dronning Maud Land, Sør Rondane Mountains, Antarctica | Black microbial mats on gravel near snow Western side of the Utsteinen Nunatak | 12 | BG11 | Fernández -Carazo et al. 2012 |
| ULC149 | *Hassallia andreassenii* AW20 | Second nunatak of Pingvinane range, Dronning Maud Land, Sør Rondane Mountains, Antarctica | Sample AW20:small stones (fallen debris?) between bigger stones on a small platform with a lot of 'black stuff' | 12 | BG110 | |
| ULC153 | *Hassallia andreassenii* PCR8 | Dronning Maud Land, Sør Rondane Mountains, Antarctica | Unknown | 18 | BG110 | |
| ULC174 | *Phormidium* | Tanngarden, | Granite | 12 | BG11 | |

| | | | | | | |
|---|---|---|---|---|---|---|
| | *lumbricale* OTC8 | Dronning Maud Land, Sør Rondane Mountains, Antarctica | outcrop on the northern side of Tanngarden, black biofilms on granitic gravel | | | |
| ULC180 | *Nostoc* sp. OTC7 | Tanngarden, Dronning Maud Land, Sør Rondane Mountains, Antarctica | Granite outcrop on the northern side of Tanngarden, black biofilms on granitic gravel | 12 | BG110 | |
| ULC186 | *Leptolyngbya* sp. FW074 | Lasne, Wallonia, Belgium | Lake Renipont water sample | 20-25 | BG11 | |
| ULC188 | *Cyanobium* sp. FW064 | Silenrieux (Eau d'Heure), Wallonia, Belgium | Lake Féronval, water sample | 20-25 | BG11 | |
| ULC191 | *Cyanobium* sp. KOTOKEL 10,1 | Unknown, Buryat Republic, Russian Federation | Freshwater from lake Kotokel | 20-25 | BG11 | |
| ULC194 | *Chroococcidiopsis* sp. PB1101 | Perlebandet, Dronning Maud Land, Sør Rondane Mountains, Antarctica | Weathered marble outcrop, NE side, exposed to wind, under patch of snow. Piece of marble. | 12 | BG11 | |
| ULC195 | *Chroococcidiopsis* sp. SV1103 | Svindlandfjellet, Dronning Maud Land, Sør Rondane Mountains, Antarctica | Big boulder, endolith on the upper North surface in cracks with very small white and black lichens | 12 | BG11 | |
| ULC197 | *Chroococcidiopsis* sp. PB1113 | Perlebandet, Dronning Maud Land, Sør Rondane Mountains, Antarctica | On the scree of garnet-biotite gneiss. In the depression of the rock, near the snow. NW ridge, lichen crust wet | 12 | BG11 | |
| ULC307 | *Phormidium autumnale* P4 | first nunatak of the Pingvinane system, westerly from Utsteinen, Dronning Maud Land, Sør Rondane Mountains, Antarctica | Sample of Prasiola in a locality with a skua nest and a lot of remains of Snow Petrels | 20-25 | BG11 | |
| ULC345 | *Leptolyngbya* sp. P.RUS1 | Moorea, French Polynesia, France | Tissue of coral Porites rus | 20-25 | ASNIII | |
| ULC346 | *Myxosarcina* sp. P.RUS2 | Moorea, French Polynesia, France | Tissue of coral Porites rus | 20-25 | ASNIII | |
| ULC367 | *Pegethrix frigida* KOVACIK ANT 2003/73 | Keller Peninsula, King Georges Island, Admiralty Bay, Antarctica | Dry epilithic crust in seepage | 12 | BG11 | Jancusova et al. 2016 |
| ULC369 | *Wilmottia murrayi* KOVACIK ANT 2003/15 | Keller Peninsula, King Georges Island, Admiralty Bay, Antarctica | Black skinny crust on moss near the Catharacta | 12 | BG11 | Jancusova et al. 2016 |

| | | | | | | |
|---|---|---|---|---|---|---|
| | | | maccormicki nest | | | |
| ULC371 | *Microcoleus attenuatus* KOVACIK ANT 2003/43 | Keller Peninsula, King Georges Island, Admiralty Bay, Antarctica | Dark green growth on remainder of whale bone near seashore | 12 | BG11 | Jancusova et al. 2016 |
| ULC373 | *Microcoleus favosus* KOVACIK ANT 2004/50 | Copacabana, King Georges Island, Admiralty Bay, Antarctica | Green soil powder near a penguin rookery side | 12 | BG11 | Jancusova et al. 2016 |
| ULC376 | *Wilmottia murrayi* KOVACIK ANT 2004/75 | Whalers Bay, Antarctic Peninsula, Deception Island, Antarctica | Mud from a littoral shallow coastal pool | 12 | BG11 | Jancusova et al. 2016 |
| ULC381 | *Geitlerinema* sp. L2 | Goma, Congo | Lac Vert, water sample | 20-25 | BG11 | |
| ULC384 | *Leptolyngbya* sp. L6 | Goma, Congo | Lac Vert, water sample | 20-25 | BG11 | |
| ULC389 | *Pseudanabaena* sp. L27 | Goma, Congo | Lac Vert, water sample | 20-25 | BG11 | |
| ULC397 | *Nostoc* sp. TG3 | Moraine slope west of the Utsteinen nunatak, Dronning Maud Land, Sør Rondane Mountains, Antarctica | Small gravel particles between the rocks, moraine | 12 | BG110 | |
| ULC401 | *Timaviella circinata* GR4$^{T}$ | Trieste, Italy | Lampenflora, on rock surface in the Giant Cave | 18 | BG11 | Sciuto et al. 2017 |
| ULC402 | *Timaviella karstica* GR13$^{T}$ | Trieste, Italy | Lamenflora growing on rock surface in the Giant Cave | 18 | BG11 | Sciuto et al. 2017 |
| ULC403 | *Phormidium autumnale* FB2 | Hverfell volcano, Sub-arctic, Iceland | Fine particles of rhyolite dust | 20-25 | BG11 | |
| ULC406 | *Tychonema bornetii* YLC011 | Cape Day, South Victoria Land, Antarctica | Microbial mat | 18 | BG11 | |
| ULC409 | *Nostoc* sp. TG10 | Utsteinen Ridge, Dronning Maud Land, Sør Rondane Mountains, Antarctica | Soil with dark green fragments | 18 | BG110 | |
| ULC415 | *Geitlerinema* sp. SHC | Gilan Province, Caspian Sea, Iran | Unknown | 18 | BG11 | |
| ULC417 | *Geitlerinema amphibium* Gor-1 | Kurdistan, Iran | BaBa Gorgor mineral water spring | 18 | BG11 | |
| ULC421 | *Nodularia spumigena* CCY9414 | Bornholm Sea, Germany | Surface water | 18 | 2/3 BG110 + 1/3 ASNI | Kopf et al. 2015 |
| ULC422 | *Thermoleptolyngbya albertanoae* ETS-08 | Padova, Euganean Thermal District, Italy | Montegrotto Terme | 20-25 | BG11 | Sciuto et al. 2016 |
| ULC424 | *Leptolyngbya ectocarpi* C86 | STARESO station, Corsica, Calvi, France | Epiphytic on Halopteris scoparia growing on a quay | 25 | ASNIII | Ohki et al. 2004; Wilmotte et al., 1988 |
| ULC426 | *Phormidium ambiguum* HK2 | Lake Winnipeg, Manitoba, Canada | Boat harbor beach, splash zone at the beach area | 12 | BG11 | |
| ULC428 | *Cyanobium* sp. Torca | Laguna Torca | Lake Torca | 20-25 | BG11 | |

| | | | | | | |
|---|---|---|---|---|---|---|
| | | National Reserve, Maule Region, Chile | | | | |
| ULC429 | *Leptolyngbya gracilis* POS1 | Bay of the scientific station STARESO (La Revellata), Corsica, Calvi, France | Epiphytic on a rhizome of Posidonia oceanica | 18 | ASNIII | |
| ULC431 | *Leptolyngbya* sp. LK1 | Lake Katinda, Uganda | Lake water | 20-25 | BG11 | |
| ULC437 | *Jaaginema subtilissimum* La Gombe 3 | Quarry of La Gombe, Esneux, Belgium | Elodea growing on rocks, 4 to 5 m depth underwater | 20-25 | BG11 | |
| ULC441 | *Phormidium papyraceum* BOTA3 | Botany Institute B22, Sart-Tilman campus, Belgium | Bloc of concrete | 18 | BG11 | |
| ULC444 | *Arthrospira platensis* Lonar | Lonar Lake, India | Alkaline Lake | 20-25 | Zarrouk | |
| ULC445 | *Arthrospira platensis* Paracas | Unknown | Unknown | 20-25 | Zarrouk | |
| ULC447 | *Nostoc* sp. TG11 | Utsteinen Ridge, Dronning Maud Land, Sør Rondane Mountains, Antarctica | Brown soil under lichen on the Utsteinen Ridge | 18 | BG110 | |
| ULC448 | *Spirulina subsalsa* BLCC-M33 | Black Point Marina, Florida, USA | Benthic | 20-25 | ASNIII | |
| ULC449 | *Leptolyngbya* sp. BLCC-M10 | Everglades National Park, Florida, USA | Benthic | 20-25 | ASNIII | |
| ULC453 | *Leptolyngbya* sp. BLCC-M28 | Jupiter Park, Florida, USA | Benthic | 20-25 | ASNIII | |
| ULC454 | *Oscillatoria* sp. BLCC-M29$^{T}$ | Black Point Marina, Florida, USA | Benthic | 20-25 | ASNIII | |
| ULC457 | *Vermifilum ionodolium* BLCC-M32 | Black Point Marina, Florida, USA | Benthic | 20-25 | ASNIII | Berthold et al., 2021b |
| ULC468 | *Cyanobium* sp. BS5-0 | Bornholm Sea, Baltic sea, Danemark | 10 m depth water | 18 | 2/3 BG11 + 1/3 ASNII | Ernst et al., 2003 |
| ULC471 | *Cyanobium* sp. BS4-0 | Bornholm Sea, Baltic sea, Danemark | 10 m depth water | 18 | 2/3 BG11 + 1/3 ASNII | Ernst et al., 2003 |
| ULC483 | *Cyanobium* sp. 6007 | Latyan dam, Iran | Surface water samples (50 cm depth) | 20-25 | BG11 | |
| ULC484 | *Leptolyngbya* sp. 6008 | Taleqan dam, Iran | Surface water samples (50 cm depth) | 20-25 | BG11 | |
| ULC500 | *Geitlerinema nematodes* W1.1.1 | Vlaams Brabant, Tervuren, Belgium | Ornemental pond | 20-25 | BG11 | |
| ULC503 | *Geitlerinema* cf. *ionicum* W1.2.2 | Vlaams Brabant, Tervuren, Belgium | Ornemental pond | 20-25 | BG11 | |
| ULC504 | *Geitlerinema* cf. *ionicum* W2.1.2 | Vlaams Brabant, Tervuren, Belgium | Ornemental pond | 20-25 | BG11 | |
| ULC508 | *Pseudoanabaena persicina* POS | Bay of the scientific station STARESO (La Revellata), Corsica, Calvi, France | Epiphytic on a rhizome of Posidonia oceanica | 18 | ASNIII | |
| ULC522 | *Neolyngbya arenicola* BLCC-M52 | Indian River Lagoon, Florida, USA | Benthic | 20-25 | BG11 marine | Lefler et al. 2021 |
| ULC525 | *Affixifilum floridanum* BLCC-M61$^{T}$ | Elliott Key, Florida, USA | Benthic | 20-25 | BG11 marine | Lefler et al. 2021 |

| | | | | | | |
|---|---|---|---|---|---|---|
| ULC529 | *Vermifilum ionodolium* BLCC-M65 | Biscayne National Park, Florida, USA | Benthic | 20-25 | BG11 marine | Berthold et al., 2021b |
| ULC530 | *Neolyngbya biscaynensis* BLCC-M69[T] | Biscayne National Park, Florida, USA | Benthic | 20-25 | BG11 marine | Lefler et al. 2021 |
| ULC535 | *Neolyngbya biscaynensis* BLCC-M95 | Biscayne National Park, Florida, USA | Benthic | 20-25 | BG11 marine | Lefler et al. 2021 |
| ULC542 | *Affixifilum granulosum* BLCC-M111 | Cannon Beach, John Pennekamp Coral Reef State Park, Florida, USA | Benthic | 20-25 | BG11 marine | Lefler et al. 2021 |
| ULC544 | *Brasilonema santannae* BLCC-T43[T] | Mid Florida Research and Education Center, Florida, USA | Aerophytic, terrestrial | 20-25 | BG11 marine | Barbosa et al. 2021 |
| ULC545 | *Brasilonema tolantongensis* BLCC-T61 | Mid Florida Research and Education Center, Florida, USA | Aerophytic, terrestrial | 20-25 | BG11 marine | Barbosa et al. 2021 |
| ULC546 | *Brasilonema santannae* BLCC-T64 | Mid Florida Research and Education Center, Florida, USA | Aerophytic, terrestrial | 20-25 | BG11 marine | Barbosa et al. 2021 |
| ULC547 | *Brasilonema octagenarum* BLCC-T71 | Mid Florida Research and Education Center, Florida, USA | Aerophytic, terrestrial | 20-25 | BG110 | Barbosa et al. 2021 |
| ULC548 | *Brasilonema fioreae* BLCC-T72[T] | Mid Florida Research and Education Center, Florida, USA | Aerophytic, terrestrial | 20-25 | BG110 | Barbosa et al. 2021 |
| ULC550 | *Brasilonema octagenarum* BLCC-T74 | Mid Florida Research and Education Center, Florida, USA | Aerophytic, terrestrial | 20-25 | BG110 | Barbosa et al. 2021 |
| ULC551 | *Brasilonema fioreae* BLCC-T83 | Mid Florida Research and Education Center, Florida, USA | Aerophytic, terrestrial | 20-25 | BG110 | Barbosa et al. 2021 |
| ULC557 | *Neolyngbya arenicola* BLCC-M60 | Biscayne National Park, Florida, USA | Benthic | 20-25 | BG11 marine | Lefler et al. 2021 |
| ULC559 | *Spirulina subsalsa* BLCC-M70 | Biscayne National Park, Florida, USA | Benthic | 20-25 | BG11 marine | |
| ULC567 | *Limnoraphis* BLCC-M92 | Guerande, Salt Marshes, France | Benthic | 20-25 | BG11 marine | |
| ULC573 | *Brasilonema wernerae* BLCC-T49[T] | Mid Florida Research and Education Center, Florida, USA | Aerophytic, terrestrial | 20-25 | BG110 | Barbosa et al. 2021 |
| ULC574 | *Brasilonema tolantongensis* BLCC-T51 | Mid Florida Research and Education Center, Florida, USA | Aerophytic, terrestrial | 20-25 | BG110 | Barbosa et al. 2021 |
| ULC575 | *Iningainema tapete* BLCC-T55[T] | Mid Florida Research and Education Center, Florida, USA | Aerophytic, terrestrial | 20-25 | BG110 | Berthold et al., 2021a |
| ULC586 | *Neolyngbya regalis* BLCC-M54 | Hollywood Beach, Florida, USA | Benthic | 20-25 | BG11 marine | Lefler et al. 2021 |
| ULC588 | *Affixifilum granulosum* BLCC-M117 | Cannon Beach, John Pennekamp Coral Reef State Park, Florida, USA | Benthic | 20-25 | BG11 marine | Lefler et al. 2021 |

| | | | | | | |
|---|---|---|---|---|---|---|
| ULC590 | *Johannesbaptistia floridana* BLCC-M67[T] | Biscayne National Park, Florida, USA | Benthic mat on coastal marine sediment | 20-25 | BG11 marine | Berthold et al., 2020 |
| ULC591 | *Parakomarekiella sesnandensis* COI00088998[T] | Coimbra, Portugal | Biodeteriorated walls of the Old Cathedral of Coimbra | 20-25 | BG110 | Soares et al., 2020 |
| ULC597 | *Leptochromothrix valpauliae* BLCC-M82[T] | Biscayne National Park, Florida, USA | Benthic | 20-25 | BG11 marine | Berthold et al., 2021b |
| ULC598 | *Leptochromothrix engenei* BLCC-M83 | Biscayne National Park, Florida, USA | Benthic | 20-25 | BG11 marine | Berthold et al., 2021b |
| ULC599 | *Ophiophycus aerugineus* BLCC-M88[T] | Biscayne National Park, Florida, USA | Benthic | 20-25 | BG11 marine | Berthold et al., 2021b |
| ULC600 | *Ophiophycus aerugineus* BLCC-M93 | Biscayne National Park, Florida, USA | Benthic | 20-25 | BG11 marine | Berthold et al., 2021b |
| ULC601 | *Ophiophycus aerugineus* BLCC-M103 | Twin Rivers Park, Port Salerno Waterfront District, Staurt. Florida, USA | Benthic | 20-25 | BG11 marine | Berthold et al., 2021b |
| ULC602 | *Aphanizomenon* sp. BL5 | Unknown, Vlaams Brabant, Brussels Hoofdstedelijk gewest, Belgium | Lake | 20-25 | 50% BG110 | |
| ULC604 | *Chroococcus* sp. waterbottle | Neupré, Belgium | Static tab water | 20-25 | BG11 | |
| ULC610 | *Planktothrix* sp. aqua2 | Neupré, Belgium | Aquarium | 20-25 | BG11 | |
| ULC611 | *Dolichospermum* sp. BL7 | Rue de pecheries,Watermaal-Bosvoorde, Brussels Hoofdstedelijk Gewest, Belgium | Pond | 20-25 | BG110 | |
| ULC718 | *Cephalothrix komarekiana* sp. Nov[T] | Pantanal da Nhecolandia, Mato Grosso do sul, Brazil | Alkaline Lake | 20-25 | BG11 | da Silva Malone et al., 2017 |

**Supplemental file 1. Information of the 152 ULC strains used in this study, including their strain number, taxonomic name, geographical origin, isolation habitat, recommended growth temperature (°C), recommended growth medium, and if applicable, the corresponding reference.** Please find the growth medium description for BG11, BG110 and ASNII in Rippka et al. (1979), and for Zarrouk in Zarrouk (1966). “50% BG110” is BG110 medium diluted twice in MilliQ water;  “BG11 marine” is composed of BG11 with an addition of 35 g/L of aquarium sea salts; “2/3 BG110 + 1/3 ASNI” contains two third of BG110 and one third of ASNIII medium; “Zarrouk+”, is prepared as Zarrouk medium, but with an addition of 197 g/l of $Na_2CO_3$.
